# Nonlinear stimulus representations in neural circuits with approximate excitatory-inhibitory balance

**DOI:** 10.1101/841684

**Authors:** Cody Baker, Vicky Zhu, Robert Rosenbaum

## Abstract

Balanced excitation and inhibition is widely observed in cortical recordings. How does this balance shape neural computations and stimulus representations? This problem is often studied using computational models of neuronal networks in a dynamically balanced state. However, these balanced network models predict a linear relationship between stimuli and population responses, in contrast to the nonlinearity of cortical computations. We show that every balanced network architecture admits some stimuli that break the balanced state and these breaks in balance push the network into a “semi-balanced state” characterized by excess inhibition to some neurons, but an absence of excess excitation. The semi-balanced state is unavoidable in networks driven by multiple stimuli, consistent with experimental data, has a direct mathematical relationship to artificial neural networks, and permits nonlinear stimulus representations and nonlinear computations.

## Introduction

An approximate balance between excitatory and inhibitory synaptic currents is widely reported in cortical recordings [1, 2, 3, 4, 5, 6]. The implications of this balance are often studied using large networks of model neurons in a dynamically stable balanced state. Despite the complexity of spike timing dynamics in these models, their population-level firing rates [7, 8, 9, 10, 11] and correlations [12, 13, 14, 15, 16, 17] in response to a given stimulus can be derived using a simple mean-field theory.

This classical mean-field theory of balanced networks has two critical shortcomings. First, it predicts a linear relationship between stimuli and population responses, in contrast to the nonlinear computations that must be performed by cortical circuits. Secondly, parameters in balanced network models must be tuned so that the firing rates predicted by the mean-field theory are non-negative. In networks with many neural populations – such as multiple neuron subtypes, neural assemblies, or tuning preferences – the proportion of parameter space for which predicted rates are non-negative becomes exponentially small. Moreover, we show that for any network architecture, there are infinitely many excitatory stimuli for which the mean-field theory predicts negative rates.

We develop a theory of semi-balanced networks that quantifies network responses when the classical balanced network state is broken. In this semi-balanced state, balance is only enforced in one direction: neurons can receive excess inhibition, but not excess excitation. Neurons receiving excess inhibition are silenced and the remaining neurons form a balanced sub-network. Unlike balanced networks, semi-balanced networks implement nonlinear computations and stimulus representations. We establish a mathematical relationship between semi-balanced networks, artificial recurrent neural networks used for machine learning [18], and threshold-linear networks [19, 20, 21, 22]. We demonstrate that balance and semi-balance are achieved on a neuron-by-neuron basis in networks with large in-degrees and homeostatic inhibitory plasticity when exposed to a time-constant stimulus [23, 24, 25], but only semi-balance is achieved in the presence of time-varying stimuli. In this setting, semi-balanced networks implement richly nonlinear stimulus representations. We demonstrate the computational power of these representations using the hand-written digit classification benchmark, MNIST.

In summary, the large in-degrees typical of cortical neurons combined with the presence of time-varying stimuli imply that local cortical circuits are in a semi-balanced state. Our analysis of this state shows a direct correspondence to artificial neural networks used in machine learning and therefore has deep implications for the computational properties of cortical circuits.

## Results

### Balanced networks implement linear stimulus representations and computations

To review balanced network theory and its limitations, we consider a recurrent network of *N* = 3 × 10^4^ randomly connected adaptive exponential integrate-and-fire (adaptive EIF) neuron models. The network is composed of two excitatory populations and one inhibitory population (80% excitatory and 20% inhibitory neurons altogether) and receives feedforward synaptic input from two external populations of Poisson processes, modeling external synaptic input (Fig. 1A). The firing rates, ***r***_*x*_ = [*r*_*x*1_ *r*_*x*2_], of the external populations form a two-dimensional stimulus space (Fig. 1B).

**Figure 1:**
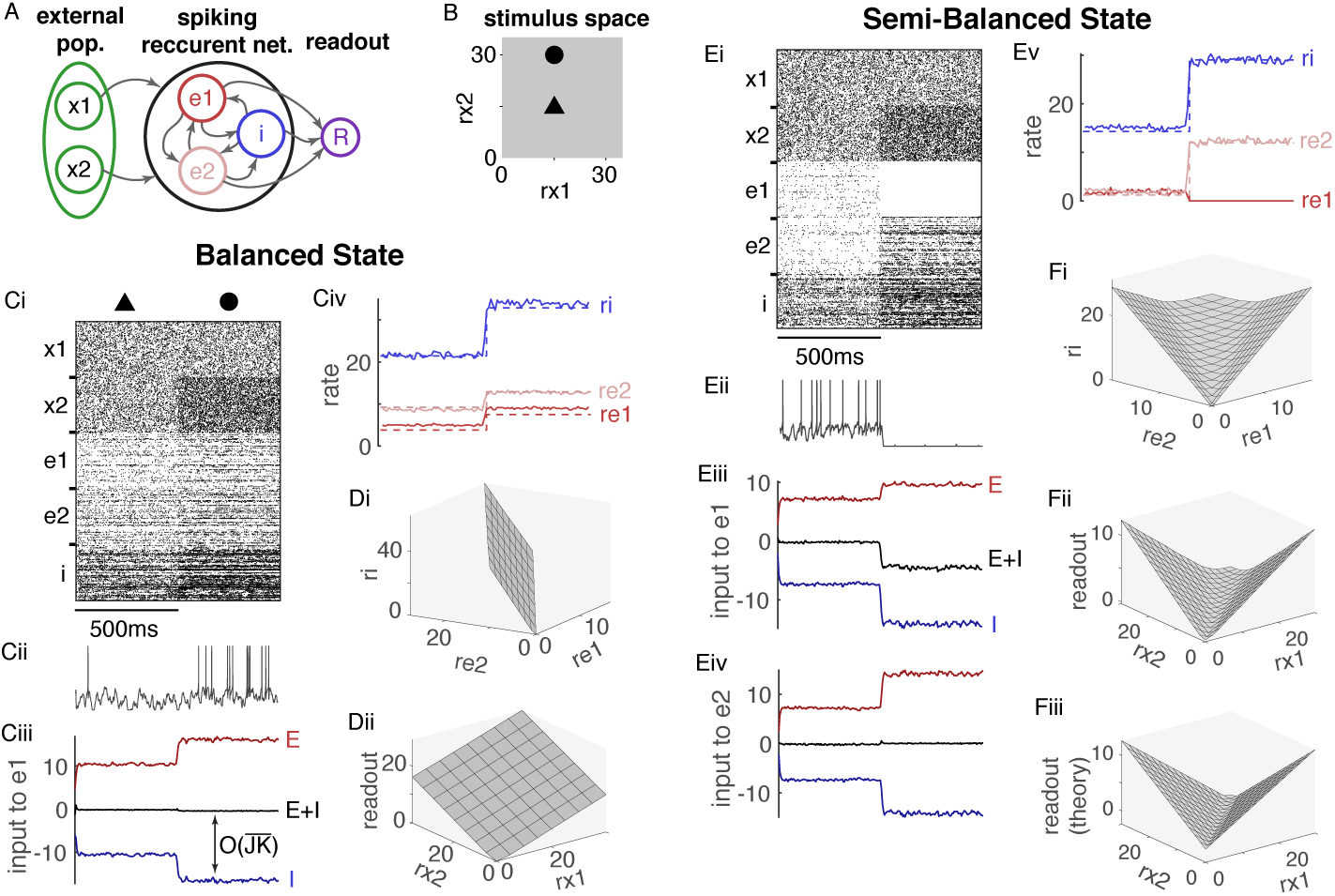
Stimulus representations are linear in the balanced state and nonlinear in the semi-balanced state. **A)** Network diagram. A recurrent spiking network of *N* = 3*×*10^4^ model neurons is composed of two excitatory populations (*e*1 and *e*2) and one inhibitory population (*i*) that receive input from two external spike train populations (*x*1 and *x*2). Recurrent network output is represented by a linear readout of firing rates (*R*). **B)** The two-dimensional space of external population firing rates represents a stimulus space. Filled triangle and circle show the two stimulus values used in Ci–iv. **Ci)** Raster plots of 200 randomly selected spike trains from each population across two stimuli. **Cii)** Membrane potential of one neuron from population *e*1. **Ciii)** Mean input current to population *e*1 from all excitatory sources (*e*1, *e*2, *x*1, and *x*2; red), from the inhibitory population (*i*; blue), and from all sources (black) showing approximate excitatory-inhibitory balance across stimuli. Mean input to *i* and *e*2 were similarly balanced. **Civ)** Firing rates of each population from simulations (solid) and predicted by Eq. (3) (dashed). **Di)** The neural manifold traced out by firing rates in each population in the recurrent network as external firing rates are varied across a square in stimulus space (0 ≤ *r*_*x*1_, *r*_*x*2_ ≤ 30). **Dii)** The readout as a function of *r*_*x*1_ and *r*_*x*2_ from the same simulation as Di. **Ei–v)** Same as Ai–iv, but dashed lines in Dv are from Eq. (4) and input to *e*2 was additionally shown. D **Fi-iii)** Same as Di-ii except the theoretical readout predicted by Eq. (4) was additionally included. All firing rates are in Hz.

Simulations of this model showed asynchronous-irregular spiking activity and excitatory-inhibitory balance (Fig. 1Ci-iii). How does connectivity between the populations determine the mapping from stimulus, ***r***_*x*_, to firing rates, ***r*** = [*r*_*e*1_ *r*_*e*2_ *r*_*i*_] in the recurrent network? Firing rate dynamics are often approximated using models of the form

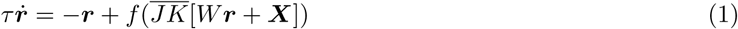

where 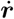 denotes the time derivative, *f* is a non-decreasing f-I curve, and *W* is the effective recurrent connectivity matrix. External input is quantified by ***X*** = *W*_*x*_*r*_*x*_. Components of *W* and *W*_*x*_ are given by 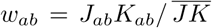 where *K*_*ab*_ is the mean number of connections from population *b* to *a* and *J*_*ab*_ is the average connection strength. The coefficient, 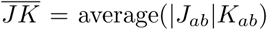, quantifies coupling strength in the network. Since 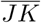 is multiplied in the equation for 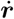 and divided in the equation for *w*_*ab*_, it does not affect dynamics, but serves as a notational tool in the calculations below, which require 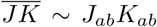 so that *w*_*ab*_ ~ 𝒪(1) even when *J*_*ab*_*K*_*ab*_ is large.

The key idea underlying balanced network theory is that 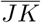 is typically large in cortical circuits because neurons receive thousands of synaptic inputs and each postsynaptic potential is moderate in magnitude. Total synaptic input,

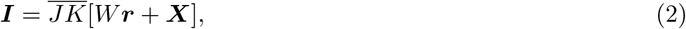

can only remain *𝒪*(1) if there is a cancellation between excitation and inhibition. In particular, to have ***I*** ∼ 𝒪(1), we must have 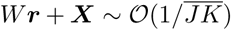 so, in the limit of large 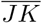, firing rates satisfy [8, 26, 11, 27]

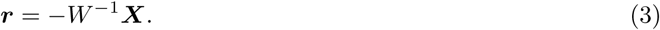

In classical balanced network theory, one considers the *N* → ∞ limit while taking 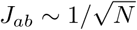 and *K*_*ab*_ ~ *N* so that 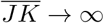 and Eq. (3) is exact in the limit [8]. Experimental evidence for this scaling has been found in cortical cultures [6]. Note that, while Eq. (1) is a heuristic approximation to spiking networks, the conclusion that Eq. (3) must be satisfied to keep ***I*** ~ 𝒪(1) as 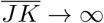 does not depend on the approximation in Eq. (1), but is implied by Eq. (2) alone and is therefore mathematically valid for spiking networks [8] for which firing rates can depend on the variance, and higher order moments of neurons’ synaptic input. Even though it is derived as a limit, Eq. (3) provides a simple approximation to firing rates in networks with finite 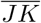. Indeed, it accurately predicted firing rates in our spiking network simulations (Fig. 1Civ, compare dashed to solid) for which 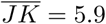 mV/Hz.

While the simplicity of Eq. (3) is appealing, its linearity reveals a critical limitation of balanced networks as models of cortical circuits: Because ***r*** depends linearly on ***X*** and ***r***_*x*_, balanced networks can only implement linear representations of stimuli and linear computations [8, 11, 27].

To demonstrate this linearity in our spiking network, we sampled a lattice of points in the two dimensional space of ***r***_*x*_ = [*r*_*x*1_ *r*_*x*2_] values and plotted the resulting neural manifold traced out in three dimensions by ***r*** = [*r*_*e*1_ *r*_*e*2_ *r*_*i*_]. The resulting manifold is approximately linear, *i.e.*, a plane (Fig. 1Di) because ***r*** depends linearly on ***X***, and therefore on ***r***_*x*_, in Eq. (3). More generally, the neural manifold is an *n*_*x*_-dimensional hyperplane in *n*-dimensional space where *n* and *n*_*x*_ are the number of populations in the recurrent and external populations respectively. In addition, any linear readout *R* = ***w · r*** is a linear function of ***r***_*x*_ and therefore also planar (Fig. 1Dii).

How do cortical circuits, which exhibit excitatory-inhibitory balance, implement nonlinear stimulus representations and computations? Below, we describe a parsimonious generalization of balanced network theory that allows for nonlinear stimulus representations by allowing excess inhibition without excess excitation.

### Semi-balanced networks implement nonlinear representations in direct correspondence to artificial neural networks of rectified linear units

Note that Eq. (3) is only valid if all elements of ***r*** it predicts are non-negative. Early work considered a single excitatory and single inhibitory population, in which case positivity of ***r*** is assured by simple inequalities satisfied in a large proportion of parameter space [8, 10]. Similarly, in the simulations described above, we constructed *W* and *W*_*x*_ so all components of ***r*** were positive for all values of *r*_*x*1_, *r*_*x*2_ > 0.

In networks with a large number of populations, conditions to assure ***r*** ≥ 0 become more complicated and the proportion of parameter space satisfying ***r*** ≥ 0 becomes exponentially small. In addition, we proved that connectivity structures, *W*, obeying Dale’s law necessarily admit some positive external inputs, ***X*** > 0, for which Eq. (3) predicts negative rates (Supplementary Materials S.1). Hence, the classical notion of excitatory-inhibitory balance cannot be assured by conditions imposed on the recurrent connectivity structure, *W*, alone, but conditions on stimuli, ***X***, are also required.

While it is possible that cortical circuits somehow restrict themselves to the subsets of parameter space that maintain a positive solution to Eq. (3) across all salient stimuli, we consider the alternative hypothesis that Eq. (3) and the balanced network theory that underlies it do not capture the full spectrum of cortical circuit dynamics.

To explore spiking network dynamics when Eq. (3) predicts negative rates, we considered the same network as above, but changed the feedforward connection probabilities so that Eq. (3) predicts positive firing rates only when *r*_*x*1_ and *r*_*x*2_ are nearly equal. When *r*_*x*2_ is much larger than *r*_*x*1_, Eq. (3) predicts negative firing rates for population *e*1, and vice versa, due to a competitive dynamic.

Simulating the network with *r*_*x*1_ = *r*_*x*2_ produces positive rates, asynchronous-irregular spiking, and excitatory-inhibitory balance (Fig. 1Ei–v, first 500ms). Increasing *r*_*x*2_ to where Eq. (3) predicts negative rates for population *e*1 causes spiking to cease in *e*1 due to an excess of inhibition (Fig. 1Ei–v, last 500ms).

Notably, however, input currents to populations *e*2 and *i* remain balanced when *e*1 is silenced (Fig. 1Eiv) so the *i* and *e*2 populations form a balanced sub-network. These simulations demonstrate a network state that is not balanced in the classical sense because one population receives excess inhibition. However,

1. no population receives excess excitation,
2. any population with excess inhibition is silenced, and
3. the remaining populations form a balanced sub-network.

Here, an excess of excitation (inhibition) in population *a* should be interpreted as 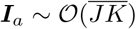 with ***I***_*a*_ > 0 (***I***_*a*_ < 0). The three conditions above can be re-written mathematically in the large 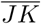 limit as two conditions,

1. [*W****r*** + ***X***]_*a*_ ≤ 0 for all populations, *a*, and
2. If [*W****r*** + ***X***]_*a*_ < 0 then ***r***_*a*_ = 0.

These conditions, along with the implicit assumption that ***r*** ≥ 0, define a generalization of the balanced state. We refer to networks satisfying these conditions as “semi-balanced” since they require that strong excitation is canceled by inhibition, but they do not require that inhibition is similarly canceled. Note that the condition [*W****r*** + ***X***]_*a*_ ≤ 0 does not mean that ***I***_*a*_ ≤ 0, but only that ***I***_*a*_ ∼ 𝒪(1) whenever ***I***_*a*_ ≥ 0 so that [*W****r*** + ***X***]_*a*_ = 0 in the large 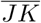 limit, *i.e.*, no excess excitation.

How are firing rates related to connectivity in semi-balanced networks? In Supplementary Materials S.2, we prove that semi-balanced networks satisfy

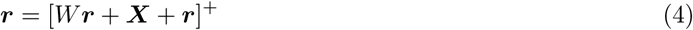

in the limit of large 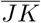 where [*x*]^+^ = max(0, *x*) is the positive part of *x*, sometimes called the rectified linear or threshold-linear function. Eq. (4) generalizes Eq. (3) to allow for excess inhibition. Even though it is derived in the limit of large 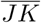, Eq. (4) provides an accurate approximation to firing rates in our spiking network simulations (Fig. 1Ev, compare dashed to solid). Note that ***r*** satisfies Eq. (4) if and only if it satisfies *q****r*** = [*W****r*** + ***X*** + *q****r***]^+^ for any *q* > 0 (see Supplementary Materials S.2 for a proof), which explains why terms with different units can be summed together in Eq. (4).

Notably, Eq. (4) represents a piecewise linear, but globally nonlinear mapping from ***X*** to ***r***. Hence, unlike balanced networks, semi-balanced networks implement nonlinear stimulus representations (Fig. 1Fi). Eq. (4) also demonstrates a direct relationship between semi-balanced networks and recurrent artificial neural networks with rectified linear activations used in machine learning [18] and their continuous-time analogues studied by Curto and others under the label “threshold-linear networks” [19, 20, 21, 22]. These networks are defined by equations of the form

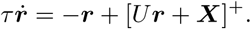

Taking *U* = *W* + *Id* where *Id* is the identity matrix establishes a one-to-one correspondence between solutions to Eq. (4) and fixed points of threshold-linear networks or recurrent artificial neural networks. Indeed, we used this correspondence to construct a semi-balanced spiking network that approximates a continuous exclusive-or (XOR) function (Fig. 1Fii–iii), which is widely known to be impossible with linear networks [18].

Previous work on threshold-linear networks shows that, despite the simplicity of Eq. (4), its solution space can be complicated [19, 20, 21, 22]: Any solution is partially specified by the subset of populations, *a*, at which ***r***_*a*_ > 0, called the “support” of the solution. There are 2^*n*^ potential supports in a network with *n* populations, there can be multiple supports that admit solutions, and these solutions can depend in complicated ways on the structure of *W* and ***X***. Hence, semi-balanced networks give rise to a rich mapping from stimuli, ***X***, to responses, ***r***.

In Supplementary Materials S.3, we proved that, under Eq. (2), the semi-balanced state is realized and Eq. (4) is satisfied only if firing rates remain moderate as 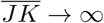. In other words, Eq. (4) and the semi-balanced state it describes are general properties of strongly and/or densely coupled networks with moderate firing rates. To the extent that cortical circuits have large 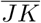 values and moderate firing rates, therefore, Eq. (4) provides an accurate approximation to cortical circuit responses. In summary, our results establish a direct mapping from biologically realistic cortical circuit models to recurrent artificial neural networks used in machine learning and to the rich mathematical theory of threshold-linear networks.

### Semi-balanced network theory is accurate across models and dynamical states

Recently, Ahmadian and Miller argued that cortical circuits may not be as tightly balanced or strongly coupled as assumed by classical balanced network theory [27]. They quantified the tightness of balance by the ratio of total synaptic input to excitatory synaptic input, *β* = |*E* + *I*|/*E* (where *E* is the mean input current from *e* and *x* combined, and *I* is the mean input from *i*). Small values of *β* imply tight balance, for example 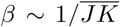 in classical balanced networks. They quantified coupling strength by the ratio of the mean to standard deviation of the excitatory synaptic current *c* = mean(*E*)/std(*E*). Strongly coupled networks have large *c*, specifically 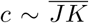. Since Eq. (4) was derived in the limit of large 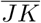, it is only guaranteed to be accurate for sufficiently large *c*, but it is not immediately clear exactly how large *c* must be for Eq. (4) to be accurate.

In our spiking network simulations, Eq. (4) was accurate across a range of stimulus values even when *β* and *c* were in the range deemed to be biologically realistic by Ahmadian and Miller (Fig. 2Ai,ii). We conclude that Eq. (4) can be a useful approximation for networks with biologically relevant levels of balance and coupling strength.

**Figure 2:**
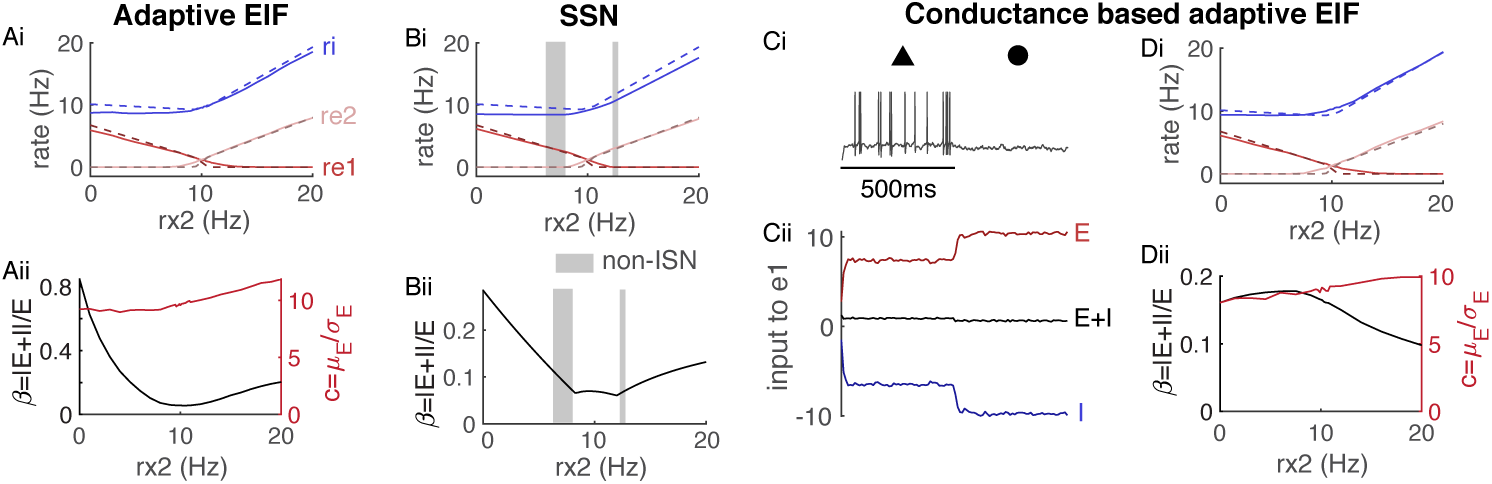
The semi-balanced approximation is accurate across models and dynamical states. **Ai)** Firing rates from simulations (solid) and Eq. (4) (dashed) as a function of *r*_*x*2_ when *r*_*x*1_ = 10Hz for the same model as in Fig. 1E-F. **Aii)** Balance ratio, *β* (black), and coupling strength coefficient, *c* (red), averaged across all neurons from the simulation in Ai. **Bi-ii)** Same as Ai and Aii, but using dynamical rate equations that implement a supralinear stabilized network (SSN). Gray shaded areas are states in which the network is not inhibitory stabilized. **Ci–ii)** Same as Fig. 1E except using a conductance-based model of synapses. **Di-ii)** Same as Ai-ii except using a conductance-based model of synapses.

We next tested the accuracy of Eq. (4) against simulations of stabilized supralinear networks (SSNs) proposed and studied by Ahmadian, Miller, and colleagues [28, 29, 27]. In particular, we simulated the three-dimensional dynamical system

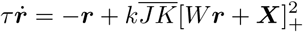

where 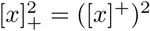 denotes the square of the positive part of *x*. Simulations of this network with parameters matched to our spiking network simulations show that the network transitioned between an inhibitory-stabilized network (ISN) state to a non-ISN state as *r*_*x*2_ varied (Fig. 2Bi), which is a defining property of SSNs. Simulations show agreement with Eq. (4), even when balance was relatively loose (Fig. 2Bi,ii).

A seemingly unrealistic property of semi-balanced networks is that the total mean synaptic current to some populations is 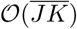 and negative (Fig. 1Eiii, black). In our simulations, this strong inhibitory input clamped the membrane potential to the lower bound we imposed at −85mV (Fig. 1Eii). The strong inhibitory current is an artifact of using a current-based model of synaptic transmission [30].

In a more realistic, conductance-based model, the magnitude of inhibitory current is limited by shunting at the inhibitory reversal potential. Repeating our simulations using a conductance-based synapse model shows similar overall trends to the current-based model (Fig. 2Ci,ii) except the mean synaptic input to population *e*1 is no longer so strongly inhibitory (Fig. 2Cii, compare to Fig. 1Eiii) and membrane potentials of *e*1 neurons still exhibit variability near the inhibitory reversal potential (Fig. 2Ci). Eq. (4) can be modified to account for conductance-based synapses (see Methods and [31, 32, 11]) and this corrected theory accurately predicted firing rates in our simulations across a range of *c* and *β* values (Fig. 2Di,ii).

### Homeostatic plasticity achieves semi-balance at single neuron resolution, producing high-dimensional nonlinear representations

So far, we have only considered firing rates and excitatory-inhibitory balance averaged over discrete neural populations. Cortical circuits implement distributed neural representations that are not always captured by homogeneous population averages [33]. Balance is realized at the level of synaptic currents to individual neurons (as opposed to currents averaged over populations) is often referred to as “detailed balance” [34, 25]. Due to the use of this term for a different purpose in Markov process theory, we instead refer to it as balance “at single-neuron resolution.”

To test for semi-balance above, we compared firing rates from simulations to those predicted by Eq. (4) (see Figs. 1Ev and 2Ai,Bi,Di). For semi-balance at single neuron resolution, or “detailed semi-balance,” Eq. (4) becomes 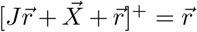. However, solving this equation is intractable for large networks because it would require searching for solutions across 2^*N*^ potential supports. Instead of comparing firing rates from simulations to those predicted by theory, we can test for semi-balance by verifying that synaptic currents to all neurons are only large in magnitude when they are negative (see Supplementary Materials S.2).

To explore balance and semi-balance at single-neuron resolution, we first considered the same randomly connected network of *N* = 3 *×* 10^4^ neurons considered above, but with only a single excitatory, inhibitory, and external population (Fig. 3A). We kept the firing rate of the external population fixed at *r*_*x*_ = 10Hz. To model a stimulus with a distributed representation, we added an extra external input perturbation that is constant in time, but randomly distributed across neurons. Specifically, the time-averaged synaptic input to each neuron was given by the *N* × 1 vector

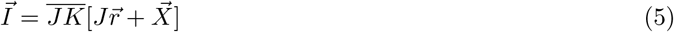

where *J* is the *N × N* recurrent connectivity matrix and 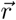 is the *N* × 1 vector of firing rates. External input is given by 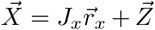 where, *J*_*x*_ and 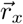 are the feedforward connectivity matrix and external rates. The distributed stimulus, 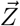, is defined by

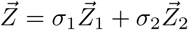

where 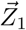 and 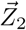 are standard normally distributed, *N* × 1 vectors. The vector, 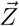, lives on a two-dimensional hyperplane in *N* -dimensional space and the plane is parameterized by *σ*_1_ and *σ*_2_. Hence, 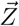 models a two-dimensional stimulus whose representation is distributed randomly across the neural population.

**Figure 3:**
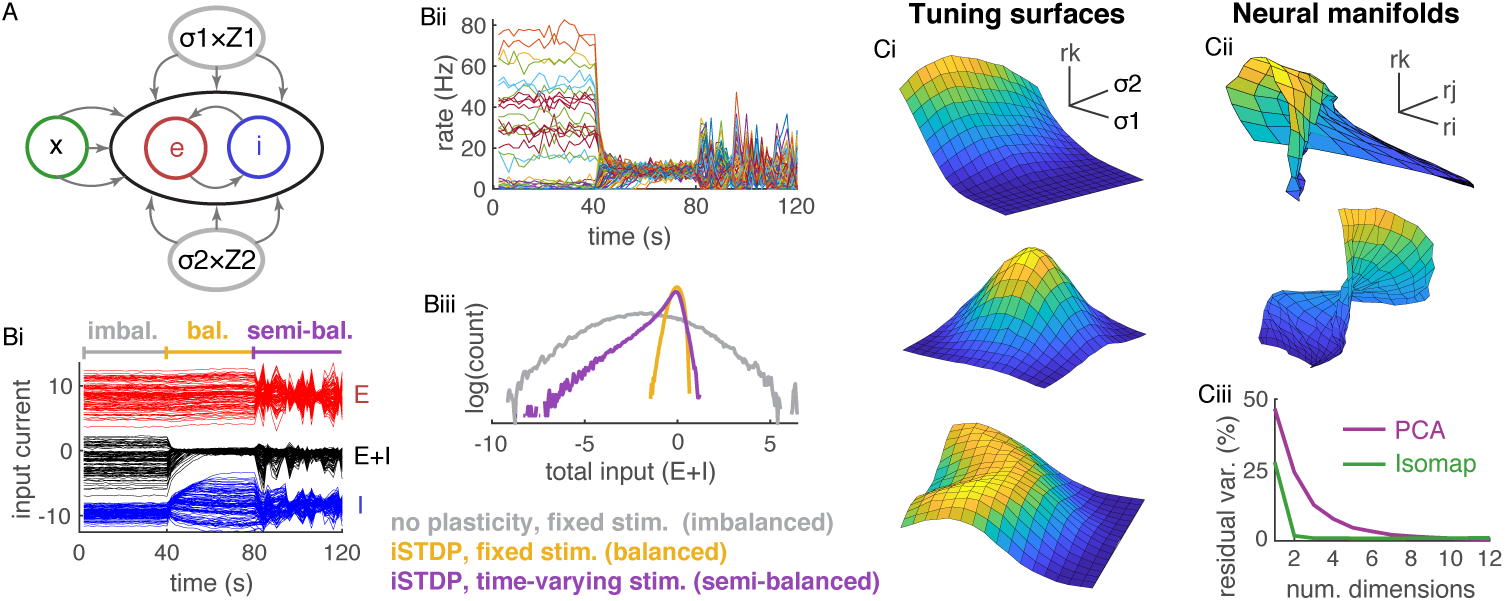
Balance, semi-balance, and neural representations at single-neuron resolution with homeostatic plasticity. **A)** Network diagram. Same as in Fig. 1A except there is just one excitatory and one external population and an additional input 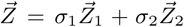. **Bi)** Excitatory (red), inhibitory (blue), and total (black) input currents to 100 randomly selected excitatory neurons averaged over 2s time bins. During the first 40s, synaptic weights and *σ*_1_ = *σ*_2_ were fixed. During the next 40s, homeostatic iSTDP was turned on and *σ*_1_ = *σ*_2_ were fixed. During the last 40s, iSTDP was on and *σ*_1_ and *σ*_2_ were selected randomly every 2s. **Bii)** Firing rates of the same 100 neurons averaged over 2s bins. **Biii)** Histograms of input currents to all excitatory neurons averaged over the first 40s (gray, imbalanced), the next 40s (yellow, balanced), and the last 40s (purple, semi-balanced). **Ci)** Firing rates of three randomly selected excitatory neurons as a function of the two stimuli, *σ*_1_ and *σ*_2_ (the neuron’s “tuning surface”) in a network pre-trained by iSTDP. **Cii)** Three neural manifolds. Specifically, the surface traced out by the firing rates of the three randomly selected neurons as *σ*_1_ and *σ*_2_ are varied. **Ciii)** Percent variance unexplained by PCA (purple) and Isomap (green) applied to all excitatory neuron firing rates from the simulation in Ci-ii. Network size was *N* = 3 *×* 10^4^ in Bi-iii and reduced to *N* = 5000 in Ci-iii to save runtime (see Methods).

Simulations show that this network does not achieve balance at single-neuron resolution: Some neurons receive excess inhibition and some receive excess excitation (Fig. 3Bi, first 40s), leading to large firing rates in some neurons (Fig. 3Biii) and a broad distribution of total input currents (Fig. 3Bii, blue). Indeed, it has been argued previously that randomly connected networks are imbalanced at single-neuron resolution when stimuli and connectivity are not co-tuned [9, 25]. This is consistent with previous results on “imbalanced amplification” in which connectivity matrices with small-magnitude eigenvalues values can break balance when external inputs are not orthogonal to the corresponding eigenvectors [11]. When *J* is large and random, it will have eigenvalues near the origin purely by chance, which can lead to imbalanced amplification if the corresponding eigenvectors are not orthogonal to 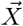.

Previous work shows that single-neuron resolution balance can be realized by a homeostatic inhibitory spike-timing dependent plasticity (iSTDP) rule [23, 25]. Indeed, when iSTDP was introduced in our simulations, balance was obtained and firing rates became more homogeneous (Fig. 3Bi-ii, second 40s) with a much narrower distribution of total input currents (Fig. 3Biii, red), at least while *σ*_1_ and *σ*_2_ were fixed.

Of course, real cortical circuits are exposed to multiple, time-varying stimuli. To simulate time-varying stimuli, we randomly selected new values of *σ*_1_ and *σ*_2_ every 2s (Fig. 3Bi-ii last 40s). This transition to a time-varying stimulus caused the total input to some neurons to become strongly inhibitory, but no neurons received excess excitation (Fig. 3Biii, yellow), indicating that the network was in a semi-balanced state at single-neuron resolution. These results show that the semi-balanced state is a natural state for cortical circuits exposed to time-varying stimuli. This is consistent with findings that inhibition dominates sensory responses in awake animals ([35], compare to dominance of inhibition in Fig. 3Bi, last 40s). Repeating these simulations in a model with conductance-based simulations shows that shunting inhibition prevents an excess inhibitory currents in the semi-balanced state, but a measure of effective excitatory and inhibitory conductances recovers the imbalanced, balanced, and semi-balanced states observed in the current-based model (Supplemental Figure S.1).

We next investigated the properties of the mapping from the two-dimensional stimulus space to the *N* -dimensional firing rate space. We sampled a uniform lattice of 17 *×* 17 = 289 points in the two-dimensional space of *σ*_1_ and *σ*_2_ values, simulated a network of size *N* = 5000, then plotted the resulting firing rates of three randomly selected neurons as a function of *σ*_1_ and *σ*_2_. The resulting surfaces appear highly nonlinear and multi-modal (Fig. 3Ci). Next, we plotted two randomly selected neural manifolds, each defined by the firing rates of three random excitatory neurons. These manifolds also appear highly nonlinear with rich structure (Fig. 3Cii). Note that there are over 10^10^ such manifolds in the network, suggesting a rich representation of the two-dimensional stimulus.

To understand how these surfaces get their shape, note that semi-balance at single-neuron resolution is realized when 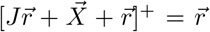 in the 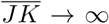 limit. This equation is piecewise linear in the sense that sufficiently small changes to 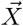 cause linear changes to 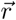. Nonlinearities occur whenever a change to 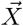 causes an individual *r*_*j*_ value to transition between zero and non-zero values, *i.e.*, at transitions between two of the 2^*N*^ “supports” (see above). The enormous number of supports in large networks (over 10^1500^ potential supports in the network from Fig. 3C) implies that nonlinearities are prevalent in the solution space and the underlying piecewise linearity is not visible in practice.

The nonlinearity of the stimulus representation is more precisely quantified by comparing the results of the dimension reduction techniques isometric feature mapping (Isomap) and principal component analysis (PCA) applied to the sampled firing rates. Both methods find a low-dimensional manifold in *N* -dimensional rate space near which the sampled rates lie. However, PCA is restricted to linear manifolds (hyperplanes) while Isomap finds nonlinear manifolds. We applied both methods to the set of all excitatory firing rates across all 289 stimuli from the simulations above.

Despite the fact that firing rates represent 289 points in a 4000-dimensional space, the points lie close to a two-dimensional manifold because they are approximately a function of the two-dimensional stimulus. Applying Isomap shows that the vast majority of the variance was explained by a two-dimensional manifold (Fig. 3Ciii, green; 1.76% residual variance at 2 dimensions). However, PCA required more than 8 dimensions to capture the same amount of variance and generally captured less variance per dimension (Fig. 3Ciii, purple). This implies that the two-dimensional neural manifold in 4000-dimensional space is nonlinear, *i.e.*, curved, so that it cannot be captured by a two-dimensional plane.

In summary, when networks are presented with time-varying stimuli, iSTDP produces a semi-balanced, but not balanced state at single neuron resolution. The mapping from stimuli to firing rates is richly nonlinear in this state. We next explore how this nonlinearity improves the computational capability of the network.

### Nonlinear representations in semi-balanced networks improve computations

To quantify the computational capabilities of our spiking networks, we used a network identical to the one from Fig. 3 except we replaced the random stimulus, 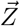, with a linear projection of pixel values from images in the MNIST data set (Fig. 4A, layer 1; see below for description of layer 2). Unlike the 2-dimensional stimuli considered previously, the images live in a 400-dimensional space (20 *×* 20 pixels).

**Figure 4:**
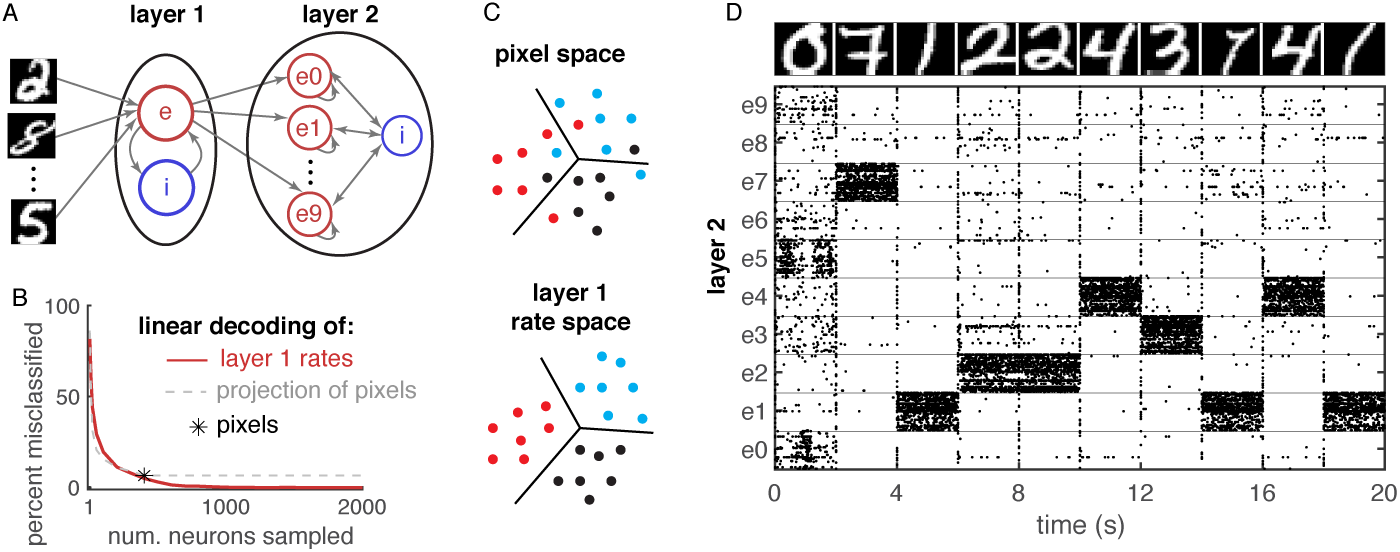
Nonlinear representations in a semi-balanced network improve computations. **A)** Network diagram. Pixel values provided external input to a semi-balanced network identical to the one in Fig. 3, representing layer 1. Layer 2 is a competitive, semi-balance network receiving external input from excitatory neurons in layer 1 with inter-laminar weights trained using a supervised Hebbian-like rule to classify the digits. **B)** Error rate (percent of 2000 images misclassified) of a linear readout of excitatory firing rates from layer 1 with readout weights optimized to classify the images, plotted as a function of the number, *n*, of neurons sampled (red). Black asterisk shows the error rate of an optimized readout of the *n* = 400 image pixels. Dashed gray shows the error rate of an optimized readout of a random projection of the pixels into *n* dimensions. The error rate of the rate readout (red curve) is zero for *n ≥* 1600. Hence, the digits are linearly separable in rate space, but not pixel space, which is only possible for nonlinear mappings. **C)** Diagram illustrating linear separability in rate space, but not pixel space. Different colors represent different digits and black lines are separating hyperplanes. **D)** Raster plot of 500 randomly selected neurons from layer 2 (50 from each population, *ek*) when images at top were provided as external input to layer 1.

We first trained inhibitory synaptic weights with iSTDP using 100 MNIST images presented for 1s each. We then presented 2000 images to the trained network and recorded the firing rates over each stimulus presentation. Applying the same Isomap and PCA analysis used above to these 2000 firing rate vectors confirms that the network implements a nonlinear representation of the images (Supplementary Figure S.2).

We wondered if the nonlinearity of this representation imparted computational advantages over a linear representation. The 10 different digits (0-9) form ten clusters of points in the 4000-dimensional space of layer 1 excitatory neuron firing rates. Similarly, the raw images represent ten clusters of points in the 400-dimensional pixel space. Are these clusters of points linearly separable?

To answer this, we trained an optimal linear readout of the 2000 firing rate vectors and found that the 10 different clusters of points in firing rate space were perfectly linearly separable. Specifically, we found a 10 *×* 4000 matrix, *W*_*r*_, such that the 10-dimensional vector, 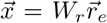, is maximized at the entry corresponding to the correct digit across all 2000 images. Here, 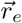 is the 4000 × 1 vector of excitatory neuron firing rates in layer 1.

For comparison, we used the same method to train an optimal linear readout of the 2000 raw MNIST images, treated as vectors in 400-dimensional pixel space. This analysis revealed that 6.6% of the images were misclassified (Fig. 4B, asterisk), implying that the digits are not linearly separable in pixel space. Hence, the digits are separable in rate space, but not in pixel space (Fig. 4C). Since the images are not linearly separable in pixel space, any linear representation of the images is not linearly separable. Hence, the separability in rate space is due to the nonlinearity of the neural representation.

We next investigated how many neurons or encoding dimensions were necessary to achieve linear separability. First, we trained an optimal linear readout on *n* randomly selected layer 1 excitatory neurons and computed the percentage of the 2000 images that were misclassified. The error decreased with *n* and perfect linear separation (zero error) was achieved for *n ≥* 1600 (Fig. 4B, red).

To compare this to pixel space, we projected each raw image randomly into *n*-dimensional space and trained a linear readout. The error of this readout for *n* ≤ 400 was similar to the error in rate space (Fig. 4B, compare gray dashed to red). However, the error in pixel space saturated to 6.6% at *n* = 400 because a linear projection of pixels into a higher dimensional space cannot improve linear separability (Fig. 4B, gray dashed curve saturates at *n* = 400).

These results demonstrate that the nonlinearity of our network improves linear discriminability of stimuli, but they do not address how well the trained linear readout performs on images that were not used in training. Moreover, the readout weights have mixed sign and do not respect Dale’s law. We next considered a downstream spiking network, layer 2, that receives synaptic input from excitatory neurons in layer 1 (Fig. 4A). Layer 2 has ten excitatory populations and one inhibitory population. Excitatory populations are coupled to themselves and bi-directionally with the inhibitory population, but do not connect to each other, producing a competitive dynamic between the excitatory populations in layer 2.

Our goal was to train feedforward weights from excitatory neurons in layer 1 to those in layer 2 that are strictly positive and encourage the *k*th excitatory population in layer 2 to be most active when layer 1 receives the digit *k* as input. We used a simple, Hebbian like learning rule in which the weight from neuron *i* in layer 1 to neuron *j* in population *ek* of layer 2 is increased when neuron *i* is active during the presentation of digit *k*. This rule is not optimal, but maintains positive weights. We applied the rule to the same 2000 images mentioned above, then tested the performance of the learned weights on 200 images not previously presented to the network. In 72.5% of these 200 test images, the network guessed the correct digit in the sense that population *ek* in layer 2 had the highest firing rate when digit *k* was presented (Fig. 4D).

## Discussion

We introduced the semi-balanced state, defined by an excess of inhibition without an excess of excitation. This state is realized naturally in networks for which the classical balanced state cannot be achieved and networks in this state implement nonlinear stimulus representations, which are not possible in classical balanced networks. We established a direct mathematical relationship between firing rates in semi-balanced networks, artificial neural networks, and the rich mathematical theory of threshold-linear networks. The semi-balanced state is realized at single-neuron resolution in networks with iSTDP, which implement high-dimensional nonlinear stimulus representations that improve the network’s computational properties.

Previous work revealed multi-stability and nonlinear transformations at the level of population averages by balanced networks with short term synaptic plasticity [36]. Future work should consider how the non-linearities introduced by short term plasticity combine with the non-linearities introduced by semi-balance. Other work studied spike timing reliability and nonlinear representations at single-neuron resolution in non-plastic networks that satisfy balance at the level of population-averages [37, 38]. Since these studies did not implement iSTDP or similar mechanisms, our results suggest that their networks were not balanced at single-neuron resolution. Hence, these studies combined with our results support the general conclusion that while networks can only perform linear computations at the resolution over which they are balanced, they can perform non-linear computations at a finer resolution. A deeper mathematical understanding of this idea is a potential topic for future work.

An alternative theory of nonlinear computations in cortical circuits is given by the theory of SSNs with power-law f-I curves [28, 29, 27]. For large 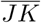, fixed point firing rates in these networks converge to the balanced fixed point, Eq. (3), when it is positive. At finite 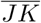, they implement nonlinearities that are not accounted for by Eq. (3). These nonlinearities are necessary to capture some experimentally observed response properties [29] and are distinct from the nonlinearities produced by semi-balance and discussed here. Indeed, fixed point firing rates in SSNs can be expanded in a series for which Eq. (3) is the first term [28]. This expansion is derived under the assumption that rates are positive, which implies that the nonlinearities produced by semi-balance are not present. Semi-balanced network theory (dashed) gives a piecewise-linear approximation of firing rates that captures the overall trends, but misses the curvature of firing rates as the stimulus changes (Fig. 2Bi, compare solid to dashed). Balanced network theory and the series expansion for SSNs are restricted to the regime in which all rates are positive. The fact that spiking networks and SSNs deviate from semi-balanced network theory in similar ways (solid and dashed differ similarly in Fig. 2Ai and Bi) suggests that SSNs can be used to refine the analysis of spiking network simulations beyond the more coarse-grained description provided by semi-balanced network theory. A key component of this analysis would be to generalize the series expansion for SSNs so that Eq. (4) is the first term instead of Eq. (3).

One limitation of our approach is that it focused on fixed point rates and did not consider their stability or the dynamics around fixed points. Previous work shows that balanced networks can exhibit spontaneous transitions between attractor states [39] which can be formed by iSTDP [23, 40]. Attractor states in those studies maintained strictly positive firing rates across populations, keeping the networks in the classical balanced state. This raises the question of whether similar attractors could arise in which some populations are silenced by excess inhibition, putting them in a semi-balanced state. Tools for studying these states could potentially be developed from the mathematical theory of threshold-linear networks [19, 20, 21, 22].

Another limitation is that, in our network trained on MNIST digits, the recurrent connections in the network were only trained via an unsupervised iSTDP rule, which is agnostic to the image labels. Hence, the recurrent network did not learn a label-dependent representation of the stimuli. Moreover, recurrent excitatory weights were not trained. Gradient-descent based learning rules for excitatory weights are easy to derive using Eq. (4) since ***r*** depends linearly on ***X*** wherever ***r*** is positive and the gradient is zero elsewhere. Future work should consider excitatory synaptic plasticity in the recurrent network and supervised learning rules for recurrent weights to learn more informative representations.

We considered violations of the balanced state arising when Eq. (3) predicts negative rates. Previous work has shown that balance can also be broken when the connectivity matrix, *W*, in Eq. (3) is singular [9, 26, 11]. While singular *W* may at first seem contrived, it has been shown that singular or nearly singular *W* arise naturally when modeling structural heterogeneity of network architectures [9, 26], optogenetic stimulation [11], or connectivity structures that depend on continuous quantities like neuron distance or orientation tuning [10, 11]. In networks with singular *W*, Eq. (4) can admit a solution even when Eq. (3) does not. Hence, semi-balanced network theory can be applied to these models for which classical balanced network theory fails. Applying semi-balanced network theory to networks with spatially continuous connectivity structure would require extending the theory of spatially extended balanced networks [41, 10, 42, 11] to account for semi-balance, *i.e.*, for spatially localized regions of neurons with quenched firing rates, which could be a fruitful direction for future work.

The semi-balanced state is defined by an excess of inhibition without a corresponding excess of excitation. This is consistent with findings that inhibition dominates cortical responses in awake animals [35]. However, it should be noted that the dominance of inhibitory synaptic currents is reduced to some extent when shunting inhibition is accounted for (see Fig. 2Cii and Supplementary Figure S.1). A more precise prediction of our model is that time-varying stimuli will silence a subset of neurons through shunting inhibition and an effective imbalance between excitatory and inhibitory conductances (see Supplementary Figure S.1 and its caption). This is consistent with evidence that visual inputs evoke shunting inhibition in cat visual cortex [43]. These predictions should be tested more precisely using *in vivo* recordings.

Recurrent spiking neural networks are notoriously difficult to train in part because the mathematical analysis of firing rates in biologically realistic recurrent spiking neural networks is largely intractable, though some approximations have been developed for some models in some parameter regimes []. Hence, gradient-based methods for firing rates in recurrent spiking networks are difficult to derive because the firing rates themselves are unknown. The piecewise linearity of firing rates in the semi-balanced state (see Eq. (4)) could greatly simplify the training of recurrent spiking networks because the gradient of the firing rate with respect to the weights can be easily computed. Future work should consider the derivation of gradient-based learning rules from Eq. (4)

Artificial recurrent neural networks for machine learning often use sigmoidal activation functions instead of the rectified linear activations typically used in feedforward networks because the unboundedness of rectified linear units make recurrent networks susceptible to instabilities and large activations [18]. However, sigmoidal activations introduce the potential for vanishing gradients that can be problematic for training [18]. Our results suggest that a homeostatic learning rule akin to an iSTDP rule could help stabilize artificial recurrent neural networks with rectified linear activations while avoiding the problem of vanishing gradients.

In summary, semi-balanced networks are more biologically parsimonious and computationally powerful than widely studied balanced network models. The foundations of semi-balanced network theory presented here open the door to several directions for further research.

## Methods

We modeled a network of *N* adaptive exponential integrate-and-fire (adaptive EIF) neurons with 0.8*N* excitatory neurons and 0.2*N* inhibitory neurons. For the current-based model used in all figures except Fig. 2B,C, the membrane potential of neuron *j* = 1, …, *N*_*a*_ in population *a* obeyed

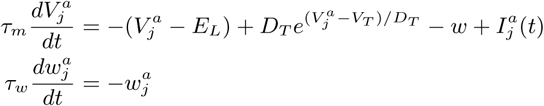

with the added condition that each time 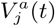 crossed *V*_*th*_ = *−*55, a spike was recorded, it was reset to *V*_*re*_ =, and 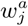 was incremented by *B* =mV. A hard lower bound was imposed at *V*_*lb*_ = *−*85mV. Other neuron parameters were *τ*_*m*_ =, *E*_*L*_ =, *D*_*T*_ =, *V*_*T*_ =, and *τ*_*w*_ =. Input was given by

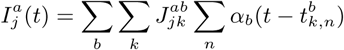

where 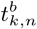 is the *n*th spike of neuron *k* in population *b* and *α_b_*(*t*) = *e^−t/τ_b_^/τ_b_H*(*t*) is an exponential postsynaptic current with *H*(*t*) the Heaviside step function. Synaptic time constants, *τ*_*b*_, were 8/4/10 ms for excitatory/inhibitory/external neurons. Synaptic weights were generated randomly and independently by

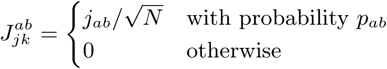

In Fig. 1C,E and Fig. 2C, external input rates were ***r***_*x*_ = [15 15]Hz for the first 500ms and ***r***_*x*_ = [15 30]Hz for the next 500ms.

In Figs. 1 and 2, postsynaptic populations were *a* = *e*1, *e*2, *i* and presynaptic populations were *b* = *e*1, *e*2, *i, x*1, *x*2 with *N*_*e*1_ = *N*_*e*2_ = 1.2 *×* 10^4^, *N*_*i*_ = 6000, and *N*_*x*1_ = *N*_*x*2_ = 3000 so that *N* = *N*_*e*1_ + *N*_*e*2_ + *N*_*i*_ = 3 *×* 10^4^. Neurons in external populations, *x*1 and *x*2, were not modeled directly, but spike times were generated as independent Poisson processes with firing rates *r*_*x*1_ and *r*_*x*2_. Connection strength coefficients were *j*_*ejek*_ = 37.5, *j*_*eji*_ = *−*225, *j*_*iek*_ = 168.75, *j*_*ii*_ = *−*375, *j*_*ejxk*_ = 2700, and *j*_*ixk*_ = 2025mV/Hz for *j, k* = 1, 2. Note that these were scaled by 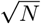 to get the actual synaptic weights as defined above. Connection probabilities in Fig. 1C,D were *p*_*e*1*e*1_ = *p*_*e*2*e*2_ = 0.15, *p*_*e*1*e*2_ = *p*_*e*2*e*1_ = 0.05, *p*_*e*1*x*1_ = 0.08, *p*_*ix*1_ = *p*_*ix*2_ = 0.12, and *p*_*ab*_ = 0.1 for all other connection probabilities. Connection probabilities in Fig. 1E,F and in Fig. 2 were the same except *p*_*e*1*x*1_ = *p*_*e*2*x*2_ = 0.15, *p*_*e*1*x*2_ = *p*_*e*2*x*1_ = 0, and *p*_*ix*1_ = *p*_*ix*2_ = 0.15.

For Fig. 2C,D, we used the model except

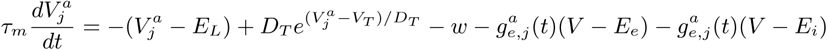

where *E*_*e*_ = 0mV, *E*_*i*_ = *−*75mV,

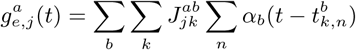

with the sum taken over excitatory presynaptic populations (*b* = *e*1, *e*2, *x*1, *x*2), and

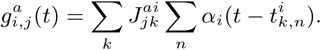

The excitatory presynaptic weights (*j*_*ae*1_, *j*_*ae*2_, *j*_*ax*1_, and *j*_*ax*2_) were the same as above, but multiplied by (*E*_*e*_ − *V*_0_) to account for the change of units. Similarly, presynaptic weights (*j*_*ai*_) were multiplied by (*E*_*i*_ − *V*_0_). We took *V*_0_ = *V*_*T*_ = *−*55mV, but the accuracy of the theory did not depend sensitively on this choice. To obtain the dashed curves in Fig. 2Di, we used Eq. (4), but with the original values of *W* (those used for the current-based model). This is equivalent to dividing the conductance-based synaptic weights by (*E*_*e*_ − *V*_0_) and (*E*_*i*_ − *V*_0_), which is the approximation produced by a mean-field theory derived in previous work [31, 32, 11].

For Fig. 2B, we solved 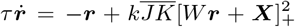 using the forward Euler method with ***r*** = [*r*_*e*1_ *r*_*e*2_ *r*_*i*_]^*T*^, ***X*** = *W*_*x*_***r***_*x*_,

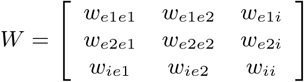

and

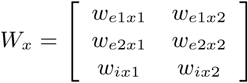

where 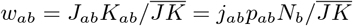. Note that 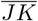 is multiplied in the differential equation and divided in the definition of *w*_*ab*_. We set *k* = 44Hz^2^/(mV)^2^ which provided a rough match to the sample f-I curves in our spiking network while still exhibiting transitions between ISN and non-ISN regimes.

For Fig. 3, the model was the same as above except there was just one excitatory, one inhibitory, and one external population with *N*_*e*_ = 0.8*N* and *N*_*i*_ = *N*_*x*_ = 0.2*N* where *N* = 3 *×* 104 in Fig. 3A,B. We reduced network size to *N* = 5 *×* 10^3^ for Fig. 3C because simulations for Fig. 3C required 289 simulations for 400s each. The long simulation time, 400s, was needed for accurate estimation of individual neuron’s firing rates at each stimulus value, which requires a longer runtime than population averaged rates. The simulation for Fig. 3C took around 54 CPU hours and run time grows quadratically with *N*, so a simulation with *N* = 3 *×* 10^4^ would have taken prohibitively long. Stimulus coefficients in Fig. 3B were set to *σ*_1_ = *σ*_2_ = 22.5mV (about 1.4 times the rheobase) for the first 80s and randomly selected from a uniform distribution on [*−*30, 30]mV for the last 40s. In Fig. 3C, *σ*_1_ and *σ*_2_ values were sampled from a uniform 17*×*17 lattice on [*−*18, 18] *×* [*−*18, 18]mV (−18mV to 18mV with a step size of 0.15 mV for each of *σ*_1_ and *σ*_2_). Connection probabilities between all populations in Fig. 3 were *p*_*ab*_ = 0.1. Initial synaptic weights were given by *j*_*ee*_ = 37.5, *j*_*ei*_ = *−*225, *j*_*ie*_ = 168.75, *j*_*ii*_ = *−*375, *j*_*ex*_ = 2700, and *j*_*ix*_ = 2025mV/Hz as above. Only inhibitory weights onto excitatory neurons (*j*_*ei*_) changed, all others were plastic.

The inhibitory plasticity rule was taken directly from previous work [23]. The variables, 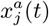, represent filtered spiking activity and are defined by 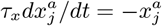 with the added condition that 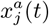 was incremented by one each time neuron *j* in population *a* = *e, i* spiked. After each spike in excitatory neuron *j*, inhibitory synaptic connections onto that neuron were updated by 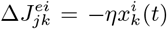 for all non-zero 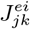. After each spike in inhibitory neuron, *k*, its outgoing synaptic connections were updated by 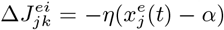. We used *τ*_*x*_ = 200ms and *α* = 2 to get a “target rate” of 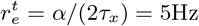.

Layer 1 in Fig. 4 was identical to the model in Fig. 3C (with *N* = 5000) except the external input was replaced by 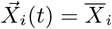 where 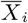 is the mean external input to inhibitory neurons in simulations with an external population (as in previous figures), so the time-varying input to inhibitory neurons was replaced by a time-constant input with the same mean. The external input to excitatory neurons was 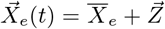 where 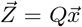 where 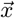 is a 400 *×* 1 vector of pixel values in the presented MNIST digit and *Q* is a *N*_*e*_ × 400 projection matrix where *N*_*e*_ = 4000. We constructed *Q* so that the *k*th pixel projected to 100 neurons, specifically to neuron indeices *j* = 100(*k −* 1) + 1 through 100*k* with strength *σ*. This corresponds to setting *Q*_*jk*_ = *σ* for 100(*k −* 1) + 1 *≤ j ≤* 100*k* and *Q*_*jk*_ = 0 otherwise. We set *σ* =?.

We first trained the inhibitory synaptic weights by presenting 100 MNIST inputs for 1 s each with iSTDP turned on. We then froze the inhibitory weights and presented an additional 2000 MNIST digits for 10 s each and saved the resulting excitatory firing rates for each digit and each excitatory neuron. Weights were frozen for this simulation because the goal is to study the (fixed) representation of digits by the trained recurrent network.

To compute the optimal readout of firing rates from Layer 1, we defined a readout *Y* = *W*_*r*_ *R*_1_ where 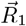 is the 4000 *×* 2000 matrix of the *N*_*e*_ = 4000 Layer 1 excitatory neuron firing rates for each of 2000 MNIST digit inputs, averaged over the 10 s that it was presented to the network. To train the 10 *×* 4000 readout matrix, *W*_*r*_, we minimized the *l*^2^ (Euclidean) norm between the 10 *×* 2000 matrix, *Y*, and the binary matrix *H* for which *H*(*m, n*) = 1 only if digit *n* = 1, …, 2000 was labeled with *m* − 1 = 0, …, 9. In other words, *H* is a matrix of one-hot vectors encoding the labeled digit. Since the *l*^2^ loss is quadratic, the minimizing *W*_*r*_ can be found explicitly. Accuracy was then computed by checking if the maximum index of *Y* was at the correct digit, *i.e.*, by taking 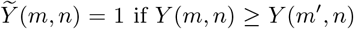 if *Y* (*m, n*) *≥ Y* (*m′*, *n*) for all *m* = 1, …, 10. As reported in Results, we obtained perfect accuracy with this procedure, *i.e.*, we obtained 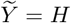 exactly. To compute the optimal readout of pixel values, represented by an asterisk in Fig. 4B, we repeated these procedures except we used the 400 *×* 1 vector of pixel values in place of the 4000 *×* 1 vector of excitatory neuron firing rates. For the red curve in Fig. 4B, we performed the same procedure, but restricted to a randomly chosen subset of the 4000 excitatory neuron firing rates (subset size indicated on the horizontal axis). For the dashed gray curve in Fig. 4B, we used a random projection, 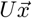, of the pixel values where 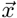 is the 400 *×* 1 vector of pixel values and *U* is a *K ×* 400 matrix with *K* being the number on the horizontal axis of the plot.

Layer 2 in Fig. 4 had *N* = 5000 neurons. The inhibitory population contained *N*_*i*_ = 1000 neurons and there were ten excitatory populations each with 400 neurons. Neurons in the same excitatory population were connected with probability *p*_*ejej*_ = 0.1 and neurons in different excitatory populations were connected with probability *p*_*ejek*_ = 0 for *j* ≠ *k*. Connection probabilities between the inhibitory population and each excitatory population were *p*_*eji*_ = *p*_*iej*_ = 0.1. Recurrent connection weights, *j*_*ab*_, were the same as for all networks considered above. Layer 2 received feedforward input from Layer 1, *i.e.*, Layer 1 served as the external input population to Layer 2.

Connectivity from Layer 1 to Layer 2 was determined as follows. We first defined a 10 *×* 400 matrix, *U*, with entries *U*_*mn*_ ≥ 0 representing connectivity from neurons in Layer 1 receiving input from pixel *k* = *n*, …, 400 to neurons in Layer 2 representing digit *m* − 1 = 0, …, 9. We trained these weights on a simulation of Layer 1 with 2000 different MNIST digit inputs. For each digit, if the digit label was *m* − 1 = 0, …, 9, we increased *U*_*mn*_ by the sum of all excitatory firing rates of neurons in Layer 1 receiving input from pixel *m*. In other words, 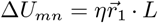 where 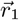 is a vector of Layer 1 firing rates and *L* = [0 … 1 … 0] is a 10 *×* 1 vector which is equal to 1 in the place of the labeled digit, *i.e.*, a one-hot vector [18]. We then normalized each column and row of *U* by its norm. This normalization makes the choice of *η* arbitrary, so we chose *η* = 1. The 4000×4000 feedforward connection matrix, *J*^21^, from excitatory neurons in Layer 1 to excitatory neurons in Layer 2 was then defined by 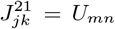 where 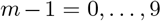 is the population to which neuron *j* = 1, …, 4000 belongs and *n* = 1, …, 400 is the pixel from which neuron *k* receives input. Inhibitory neurons in Layer 2 did not receive feedforward synaptic input, only recurrent input. Since excitatory neurons in Layer 2 are only connected to other excitatory neurons within their population, but all excitatory populations connect reciprocally to the inhibitory population, this creates a winner-take-all dynamic in which the excitatory population with the strongest external input spikes at an elevated rate and suppresses other excitatory populations. Combined with the supervised Hebbian plasticity rule, this creates a dynamic where the network learns to activate population *em* when an image is presented that is similar to training images that were labeled with digit *m*. Fig. 4D and the accuracy reported in Results reflects spiking activity in Layer 2 after training of the feedforward weights is turned off.

Matlab code to produce all figures will be included with a revised submission. Until that time, code may be requested from the corresponding author via email.

## Acknowledgments

This work was supported by NSF grants DMS-1654268 and Neuronex DBI-1707400.

## Notes

#### Summary of Updates

Minor changes to Results. Added Supplementary Materials.

