## Supplementary Materials for "Nonlinear stimulus representations in neural circuits with approximate excitatory-inhibitory balance"

### Supplementary Materials for
### Nonlinear representations in neural circuits with approximate excitatory-inhibitory balance

Cody Baker, Vicky Zhu, and Robert Rosenbaum

#### S.1 Proof that all connection matrices admit excitatory stimuli that break the classical balanced state.

Here, we prove that all connection matrices, $W$, satisfying Dale's law admit some $\boldsymbol{X}$ with positive entries for which some firing rates given by Eq. (3) in the main text are negative. The theorem relies on the presence at least one excitatory population in the network.

**Theorem 1.** *Suppose $W$ is a real, non-singular $n \times n$ matrix for which each column is either non-negative or non-positive (Dale's law), each column has at least one non-zero element, and there is at least one positive entry in the matrix. Then there exists an $n \times 1$ vector, $\boldsymbol{X}$, with strictly positive entries ($\boldsymbol{X}_j > 0$ for all $j$) for which the $n \times 1$ vector defined by $\boldsymbol{r} = -W^{-1}\boldsymbol{X}$ has at least one negative entry ($\boldsymbol{r}_j < 0$ for some $j$).*

*Proof.* Without loss of generality, we can rearrange columns to write $W$ with the non-negative columns first and the non-positive ones next,

$$W = \begin{bmatrix} + & + & \cdots & - & - \\ + & + & \cdots & - & - \\ \cdots & & & & \\ + & + & \cdots & - & - \end{bmatrix}$$

where each $+$ is an element that is $\geq 0$ and each $-$ is $\leq 0$. Now define an $n \times 1$ column vector

$$v = \begin{bmatrix} - \\ - \\ \cdots \\ + \\ + \end{bmatrix}$$

where each $-$ is a negative number, each $+$ is a positive number, there are the same number $-$ entries in $v$ as there are $+$ columns in $W$, and the same number of $+$ entries in $v$ as $-$ entries in

$W$. Finally, define

$$\boldsymbol{X} = -W\boldsymbol{v}$$

$$= -\begin{bmatrix} + & + & \cdots & - & - \\ + & + & \cdots & - & - \\ \cdots & & & & \\ + & + & \cdots & - & - \end{bmatrix} \begin{bmatrix} - \\ - \\ \cdots \\ + \\ + \end{bmatrix}$$

$$= \begin{bmatrix} + \\ + \\ \cdots \\ + \\ + \end{bmatrix}$$

In the last expression, each + is a positive number. Note that elements of $\boldsymbol{X}$ cannot be zero because of our assumption that each column of $W$ has at least one non-zero entry.

Now define, $\boldsymbol{r} = -W^{-1}\boldsymbol{X}$ and we must show that $\boldsymbol{r}$ has at least one negative entry. Compute

$$\boldsymbol{r} = -W^{-1}\boldsymbol{X} = -W^{-1}W\boldsymbol{v} = \boldsymbol{v}.$$

Therefore, $\boldsymbol{r}$ has at least one negative entry under our assumption that $W$ has at least one column with non-negative entries.

□

Note that our proof actually gives infinitely many $\boldsymbol{X}$ that satisfy the theorem, one for each $\boldsymbol{v}$ having the sign pattern defined in the proof. Moreover, there may exist additional $\boldsymbol{X}$ that are different from the ones generated by our proof.

#### S.2 Derivation and analysis of Eq. (4) from the main text.

We now prove that Eq. (4), which specifies firing rates in the semi-balanced state is equivalent to the two conditions preceding it, which define the semi-balanced state.

**Theorem 2.** *ose $W$ is an $n \times n$ matrix and $\boldsymbol{X}$ an $n \times 1$ vector. An $n \times 1$ vector, $\boldsymbol{r}$, satisfies*

*A)* $[W\boldsymbol{r} + \boldsymbol{X} + \boldsymbol{r}]^+ = \boldsymbol{r}$

*if and only if it satisfies the following three conditions at every index $a = 1, \ldots, n$:*

1. $[W\boldsymbol{r} + \boldsymbol{X}]_a \leq 0$.
2. *If* $[W\boldsymbol{r} + \boldsymbol{X}]_a < 0$ *then* $\boldsymbol{r}_a = 0$
3. $\boldsymbol{r}_a \geq 0$

*Proof.* We first show that $A$ implies conditions 1–3. Assume $\boldsymbol{r}$ satisfies $A$ and consider some index, $a$. We need to show that 1–3 are all satisfied at $a$. Condition 3 is satisfied because $\boldsymbol{r}_a = [\cdots]^+ \geq 0$.

We still need to prove that conditions 1–2 are satisfied. Note that we either have $\boldsymbol{r}_a = 0$ or $\boldsymbol{r}_a > 0$. First consider the case that $\boldsymbol{r}_a = 0$. Then 2 is satisfied automatically and we only need to prove 1. If $\boldsymbol{r}_a = 0$ then, by A, $[W\boldsymbol{r} + \boldsymbol{X}]_a^+ = \boldsymbol{r}_a = 0$ which implies that $[W\boldsymbol{r} + \boldsymbol{X}] \leq 0$. Now we must consider the case $\boldsymbol{r}_a > 0$. By A, $[W\boldsymbol{r} + \boldsymbol{X} + \boldsymbol{r}]_a^+ = \boldsymbol{r}_a > 0$, so the ReLu is evaluated at its positive part and we can conclude that $\boldsymbol{r}_a = [W\boldsymbol{r} + \boldsymbol{X} + \boldsymbol{r}]_a = [W\boldsymbol{r} + \boldsymbol{X}]_a + \boldsymbol{r}_a$. Cancelling the two $\boldsymbol{r}_a$ terms implies that $[W\boldsymbol{r} + \boldsymbol{X}]_a = 0$. Hence, 1 and 2 are both satisfied. This concludes the proof that A implies 1–3.

Now we must prove that 1–3 implies $A$. We therefore assume 1–3 and derive $A$ at each index, $a$. By 3, we must have $\boldsymbol{r}_a = 0$ or $\boldsymbol{r}_a > 0$. First assume $\boldsymbol{r}_a = 0$. Then $[W\boldsymbol{r}+\boldsymbol{X}+\boldsymbol{r}]_a^+ = [W\boldsymbol{r}+\boldsymbol{X}]_a^+ = 0$ where the last step follows from our assumption of 1. Therefore, $[W\boldsymbol{r} + \boldsymbol{X} + \boldsymbol{r}]_a^+ = \boldsymbol{r}_a = 0$. Now assume $\boldsymbol{r}_a > 0$. Then, by 1 and 2 combined, we must have $[W\boldsymbol{r} + \boldsymbol{X}]_a = 0$. Therefore, $[W\boldsymbol{r} + \boldsymbol{X} + \boldsymbol{r}]_a^+ = [\boldsymbol{r}_a]^+ = \boldsymbol{r}_a$ since $\boldsymbol{r}_a > 0$. This completes our proof.

□

Note that the condition $\boldsymbol{r}_a \geq 0$ was not explicitly included in the results because it was implicitly assumed. In the first half of our proof, we concluded that $[W\boldsymbol{r} + \boldsymbol{X}]_a = 0$ wherever $\boldsymbol{r}_a > 0$. This implies that balance is maintained at each population that has a non-zero firing rate, *i.e.*, that the populations with non-zero rates form a balanced sub-network.

The equation $[W\boldsymbol{r}+\boldsymbol{X}+\boldsymbol{r}]^+ = \boldsymbol{r}$ at first appears awkward because it sums terms with potentially different dimensions: $\boldsymbol{r}$ has dimension 1/time (*e.g.*, units Hz) while $W\boldsymbol{r}$ and $\boldsymbol{X}$ have dimensions of the neuron model's input current (measured in mV in our model since we normalized by the leak conductance, see Methods). The following theorem clarifies that this combination of dimensions is consistent because one can introduce a scaling factor without changing the solution space

**Theorem 3.** *Let $W$ be an $n \times n$ matrix and let $\boldsymbol{X}$ and $\boldsymbol{r}$ be $n \times 1$ vectors. The equation*

$$[W\boldsymbol{r} + \boldsymbol{X} + \boldsymbol{r}]^+ = \boldsymbol{r} \tag{S.1}$$

*is satisfied if and only if the equation*

$$[W\boldsymbol{r} + \boldsymbol{X} + c\boldsymbol{r}]^+ = c\boldsymbol{r} \tag{S.2}$$

*is satisfied for every $c > 0$.*

*Proof.* We first prove that Eq. (S.1) implies Eq. (S.2). Assume Eq. (S.1) is true. Let $a$ be some index. Either $\boldsymbol{r}_a = 0$ or $\boldsymbol{r}_a > 0$. First assume $\boldsymbol{r}_a = 0$. Then $[W\boldsymbol{r} + \boldsymbol{X}]_a \leq 0$ and $c\boldsymbol{r}_a = 0$. Therefore $[W\boldsymbol{r} + \boldsymbol{X} + c\boldsymbol{r}]_a^+ = [W\boldsymbol{r} + \boldsymbol{X}]_a^+ = 0 = c\boldsymbol{r}_a$. Now assume $\boldsymbol{r}_a > 0$. Then $c\boldsymbol{r}_a > 0$ and, as discussed above, we must have $[W\boldsymbol{r} + \boldsymbol{X}]_a = 0$. Therefore $[W\boldsymbol{r} + \boldsymbol{X} + c\boldsymbol{r}]_a^+ = [c\boldsymbol{r}_a]^+ = c\boldsymbol{r}_a$. This concludes our proof that Eq. (S.1) implies Eq. (S.2).

We must now prove that Eq. (S.2) implies Eq. (S.1). This is trivial because we can simply take $c = 1$. □

#### S.3 Proof that the semi-balanced state is equivalent to $\boldsymbol{r} \sim \mathcal{O}(1)$.

We now prove that for firing rate models, the semi-balanced state is realized if and only if $\boldsymbol{r} \sim \mathcal{O}(1)$ as $\overline{JK} \to \infty$. The proof relies on some reasonable assumptions on the f-I curve, *i.e.*, the function $\boldsymbol{r} = f(\boldsymbol{I})$.

**Theorem 4.** *Suppose $W$ is a fixed $n \times n$ matrix and $\boldsymbol{X}$ a fixed $n \times 1$ vector. Assume that $\boldsymbol{r}$ and $\boldsymbol{I}$ are $n \times 1$ vectors that depend on $\overline{JK}$ with*

$$\boldsymbol{I} = \overline{JK}[W\boldsymbol{r} + \boldsymbol{X}]$$

*and*

$$\boldsymbol{r} = f(\boldsymbol{I})$$

*for all sufficiently large values of $\overline{JK} > 0$. Also assume that $f(x)$ is a non-negative, non-decreasing function for which $\lim_{x\to\infty} f(x) = M$, and $\lim_{x\to-\infty} f(x) = 0$. Here, $M$ can be finite in the case of a saturating or sigmoidal f-I curve, or $M = \infty$ in the case of an f-I curve that does not saturate. If*

$$\boldsymbol{r}^{\infty} = \lim_{\overline{JK}\to\infty} \boldsymbol{r}$$

*exists and $r_a^{\infty} < M$ for all $a = 1, \ldots, n$ then*

$$[W\boldsymbol{r}^{\infty} + \boldsymbol{X} + \boldsymbol{r}^{\infty}]^{+} = \boldsymbol{r}^{\infty}. \tag{S.3}$$

*Proof.* Assume $\boldsymbol{r}^{\infty} = \lim_{\overline{JK}\to\infty} \boldsymbol{r}$ exists and is finite. Then we need to show that it satisfies Eq. (S.3). Specifically, for each index, $a = 1, \ldots, n$, we need to show that

$$[[W\boldsymbol{r}^{\infty} + \boldsymbol{X}]_a + \boldsymbol{r}_a^{\infty}]^{+} = \boldsymbol{r}_a^{\infty}$$

where $[W\boldsymbol{r}^{\infty} + \boldsymbol{X}]_a$ is the $a$th index of $W\boldsymbol{r}^{\infty} + \boldsymbol{X}$. Let $a \in \{1, \ldots, n\}$ be an arbitrary index and define

$$c = \lim_{\overline{JK}\to\infty} \frac{\boldsymbol{I}_a}{\overline{JK}}.$$

Note that

$$c = \lim_{\overline{JK}\to\infty} [W\boldsymbol{r} + \boldsymbol{X}]_a = [W\boldsymbol{r}^{\infty} + \boldsymbol{X}]_a$$

exists and is finite by assumption.

We first argue that $c \leq 0$. To show this, we will assume that $c > 0$ and prove a contradiction. If $c > 0$ then

$$\lim_{\overline{JK}\to\infty} \boldsymbol{I}_a = \lim_{\overline{JK}\to\infty} \overline{JK}c = \infty$$

and therefore

$$\boldsymbol{r}_a^{\infty} = \lim_{\overline{JK}\to\infty} f(\boldsymbol{I}_a) = M$$

which contradicts our assumption that $\boldsymbol{r}_a^{\infty} < M$ for all $M$. We may conclude that $c \leq 0$. We now break the proof into two cases: $c = 0$ and $c < 0$.

*Case 1: $c = 0$.*
We have $c = [W\boldsymbol{r}^{\infty} + \boldsymbol{X}]_a = 0$, so

$$[[W\boldsymbol{r}^{\infty} + \boldsymbol{X}]_a + \boldsymbol{r}_a^{\infty}]^{+} = [\boldsymbol{r}_a^{\infty}]^{+}$$

but $\boldsymbol{r}_a^{\infty} \geq 0$ at all indices, $a$, because $\boldsymbol{r} = f(\boldsymbol{I}) \geq 0$ at all $\overline{JK}$ and $\boldsymbol{r}^{\infty} = \lim_{\overline{JK}\to\infty} \boldsymbol{r}$. Therefore,

$$[[W\boldsymbol{r}^{\infty} + \boldsymbol{X}]_a + \boldsymbol{r}_a^{\infty}]^{+} = [\boldsymbol{r}_a^{\infty}]^{+} = \boldsymbol{r}_a^{\infty}.$$

This completes Case 1.

*Case 2:* $c < 0$.
We have $c = [W\boldsymbol{r}^\infty + \boldsymbol{X}]_a < 0$, so

$$\lim_{\overline{JK} \to \infty} \boldsymbol{I}_a = \lim_{\overline{JK} \to \infty} \overline{JK} c = -\infty.$$

Therefore,

$$\boldsymbol{r}_a^\infty = \lim_{\overline{JK} \to \infty} f(\boldsymbol{I}_a) = \lim_{\boldsymbol{I}_a \to -\infty} f(\boldsymbol{I}_a) = 0.$$

As a result,

$$[[W\boldsymbol{r}^\infty + \boldsymbol{X}]_a + \boldsymbol{r}_a^\infty]^+ = [[W\boldsymbol{r}^\infty + \boldsymbol{X}]_a]^+ = 0 = \boldsymbol{r}_a^\infty.$$

because $[W\boldsymbol{r}^\infty + \boldsymbol{X}]_a = c < 0$ and $\boldsymbol{r}_a^\infty = 0$. This completes Case 2.

□

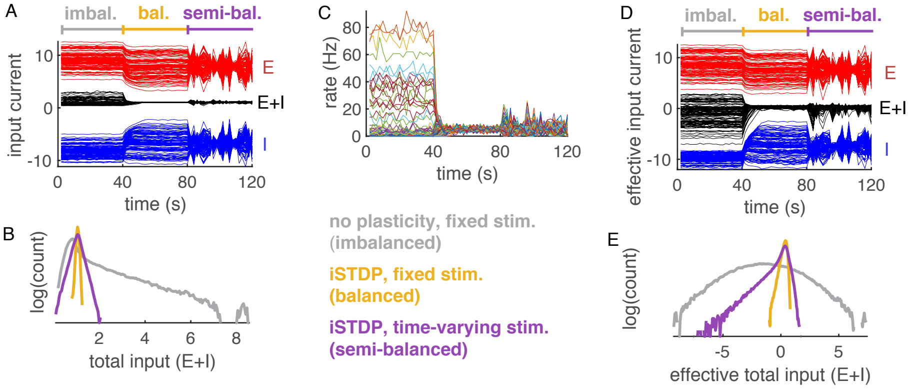


Supplementary Figure S.1: **Balance and semi-balance at single-neuron resolution in a model with conductance-based synapses. A–C)** Same as Fig. 3Bi–iii in the main text except that a conductance-based model was used for synapses. Synaptic currents from population $a$ were measured by $-g_a(t)(V(t)-E_a)$. **D–E)** Same as A–B except "effective" synaptic currents were measured by $I_a(t) = -g_a(t)(V_0 - E_a)$ where we chose $V_0 = -55$mV, but results did not depend sensitively on the choice of $V_0$. This defines a notion of effective balance and semi-balance in terms of a balance or semi-balance between the effective currents, instead of actual currents. Effective semi-balance and dominance of effective inhibition is an experimentally testable prediction of our model.

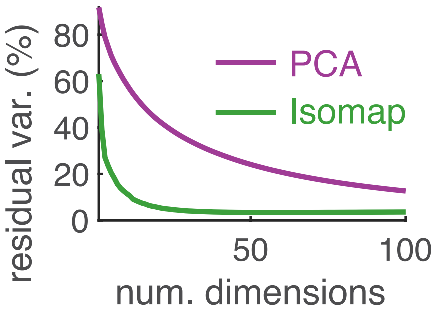


Supplementary Figure S.2: **Dimensionality of layer 1 firing rates in the model from Figure 4 of the main text.** Same as Fig. 3Ciii in the main text except IsoMap and PCA were applied to firing rates of layer 1 neurons from the model in Fig. 4 of the main text.
